# *Wolbachia* and virus alter the host transcriptome at the interface of nucleotide metabolism pathways

**DOI:** 10.1101/2020.06.18.160317

**Authors:** Amelia RI Lindsey, Tamanash Bhattacharya, Richard W Hardy, Irene LG Newton

## Abstract

*Wolbachia* is a maternally transmitted bacterium that manipulates arthropod and nematode biology in myriad ways. The *Wolbachia* strain colonizing *Drosophila melanogaster* creates sperm-egg incompatibilities and protects its host against RNA viruses, making it a promising tool for vector control. Despite successful trials using *Wolbachia*-transfected mosquitoes for Dengue control, knowledge of how *Wolbachia* and viruses jointly affect insect biology remains limited. Using the *Drosophila* model, transcriptomics and gene expression network analyses revealed pathways with altered expression and splicing due to *Wolbachia* colonization and virus infection. Included are metabolic pathways previously unknown to be important for *Wolbachia*-host interactions. Additionally, *Wolbachia*-colonized flies exhibit a dampened transcriptomic response to virus infection, consistent with early blocking of virus replication. Finally, using *Drosophila* genetics, we show *Wolbachia* and expression of nucleotide metabolism genes have interactive effects on virus replication. Understanding the mechanisms of pathogen blocking will contribute to the effective development of *Wolbachia*-mediated vector control programs.

## INTRODUCTION

*Wolbachia* is an alphaproteobacterium that establishes intracellular infections within arthropod and nematode hosts. *Wolbachia* is famous for inducing reproductive manipulations of arthropods in order to facilitate maternal transmission and spread throughout a population. In many cases, this reproductive manipulation is linked to the ability to protect the same host from secondary infections with pathogens, especially RNA viruses (Lindsey et al., 2018a). The *Wolbachia* strain infecting *Drosophila melanogaster* (*w*Mel) both induces sperm-egg incompatibilities (known as cytoplasmic incompatibility, or CI), and blocks pathogens (Teixeira et al., 2008). These phenotypes have made the *w*Mel *Wolbachia* strain highly desirable for use in vector control programs. Indeed, *Aedes aegypti* mosquitoes transfected with the *w*Mel *Wolbachia* strain form the basis of many ongoing vector control programs aimed at reducing the impact of vector-borne diseases such as Dengue and chikungunya (Hoffmann et al., 2014; Hoffmann et al., 2011a; Hoffmann et al., 2015; Walker et al., 2011).

Despite the utility of *Wolbachia* in controlling vector populations and vector borne pathogens, our understanding of the *Wolbachia*-host relationship remains limited. The pathogen blocking phenotype of *w*Mel is consistently recovered across many host species and pathogen challenges to which it has been introduced (Blagrove et al., 2013; Blagrove et al., 2012; Hughes et al., 2011a; Kambris et al., 2010; Rainey et al., 2016; Walker et al., 2011). Studies point to viruses being blocked early in infection as a result of host cell physiology that has been altered by *Wolbachia’s* presence (Bhattacharya et al., 2020; Bhattacharya et al., 2017; Geoghegan et al., 2017; Lindsey et al., 2018a; Rainey et al., 2016). However, *Wolbachia*’s effect on different hosts manifests in different ways at the cellular level, including perturbations of cholesterol availability, differential expression of host proteins, induction of the RNAi pathway, and induction of immune pathways via reactive oxygen stress (Bhattacharya et al., 2017; Geoghegan et al., 2017; Pan et al., 2011; Terradas et al., 2017; Xi et al., 2008b; Zhang et al., 2013). While these differences in host cellular environment have all been implicated in pathogen blocking, none can completely explain the phenotype across host-*Wolbachia* combinations. It is easy to imagine that *Wolbachia* would have very different effects on the intracellular environment of native and non-native hosts, which we previously reviewed in detail (Lindsey et al., 2018a).

While it is well understood that *Wolbachia* colonization results in the differential expression of host genes, it is incredibly surprising that until now this has not been investigated in *Drosophila melanogaster*, comparing *Wolbachia*-colonized and *Wolbachia*-free whole animals. Previous studies have investigated: A) immune gene expression via qRT-PCR (Bhattacharya et al., 2017), B) *Drosophila* cell lines with and without *w*Mel (Rainey et al., 2016; Xi et al., 2008a), C) whole-animal RNA-Seq in other organisms (including mosquitoes (Caragata et al., 2017; Hughes et al., 2011b; Rancès et al., 2012), nematodes (Bennuru et al., 2016), leafhoppers (Asgharian et al., 2014), and parasitoid wasps (Kremer et al., 2012)), and D) RNA-Seq of *Drosophila* and *w*Mel across fly development (but without a comparison to flies without *Wolbachia*) (Gutzwiller et al., 2015).

*Drosophila melanogaster* is the native host for *w*Mel, representing a stable host-microbe relationship (Moran et al., 2008), and the organismal context in which the pathogen blocking phenotype of *w*Mel evolved. While *Drosophila* is not a native vector for arboviruses, *Wolbachia* does significantly reduce replication of arboviruses such as Sindbis (SINV) in *Drosophila melanogaster* (Bhattacharya et al., 2017). The genetic tools available for both *Drosophila* and the type alphavirus SINV are useful for fundamental explorations of the mechanisms of intracellular infections and determinants of virus infectivity (Avadhanula et al., 2009). Ultimately, understanding mechanisms of pathogen blocking and their evolution will facilitate the long-term success of *Wolbachia*-mediated vector control. Below we present a comprehensive RNA-Seq analysis of the effect of *Wolbachia* colonization, SINV infection, and their interactive effects in the *D. melanogaster* host.

## RESULTS

### *Wolbachia* colonization and virus infection globally affect fly transcription

We generated ~1.56 billion reads with a mean quality score of 34.21 across 48 libraries to assess the impact of virus infection and *Wolbachia* colonization on fly gene expression, and of virus infection on *Wolbachia* gene expression. On average we generated 32.5 million reads per library. We detected no significant contamination in our libraries: libraries derived from *Wolbachia*-free flies had few reads mapping to the *Wolbachia* genome, and they were likely from the microbiome as they only mapped to conserved portions of rRNA genes and had perfect BLAST hits to genera such as *Lactobacillus* and *Acetobacter*, which are core components of the *Drosophila melanogaster* gut microbiome (Broderick and Lemaitre, 2012). Similarly, libraries derived from PBS-injected flies had a small proportion of reads map to the SINV genome, but these were only partial read matches that aligned to the SINV poly-A tail, and not the open reading frames of the virus. Multidimensional scaling (MDS) plots of the global similarity in fly gene expression revealed clustering of samples based on their *Wolbachia* colonization status, SINV infection status, and time post injection (Figure 1, Supplemental Table S5). The first dimension of the MDS plots separated samples by time, highlighted by the arrows overlaid on Figure 1A. Indeed, flies that were injected with SINV have very different trajectories of gene expression than did the flies that were injected with PBS alone. Flies injected with PBS do show changes in gene expression across the duration of the experiment, and this is likely due to the recovery from injection, which is distinct from the changes in gene expression experienced by flies injected with SINV. The second dimension of the MDS plots primarily separated samples based on *Wolbachia* colonization, whereas dimension three separated samples based on virus infection (Figure 1B). Three-dimensional representation of the MDS analysis showed the distinct clustering of samples based on their unique combination of *Wolbachia*-SINV-time (Figure 1C). In contrast to the fly gene expression data, *Wolbachia* gene expression did not cluster based on virus infection status (Supplementary Figure S1), indicating *Wolbachia* did not respond to SINV infection, as has been shown previously (Rainey et al., 2016).

**Figure 1.**
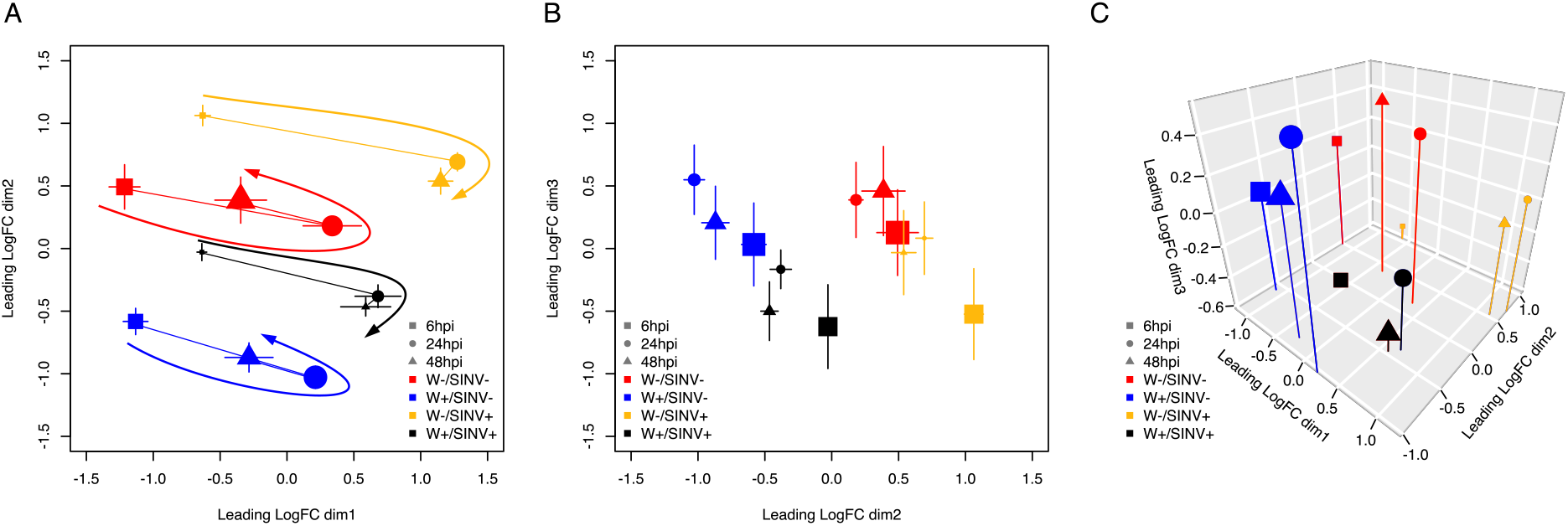
Global transcriptomic response of *Drosophila* to *Wolbachia* colonization and SINV infection. MDS plots showing similarity of total gene expression of all samples across three dimensions. Biological replicates were averaged to show their center of gravity. (A) Similarity of samples across dimension-1 and −2. The size of points indicates how close (larger points) or far away (smaller points) they are along dimension-3, which comes in and out of the page. Lines connect timepoints within a *Wolbachia* (W)/SINV combination, and arrows show the trajectory of gene expression across time. (B) Dimension-2 and −3 show a clustering of samples based on SINV-infection and *Wolbachia*-colonization. The size of points indicates how close (larger points) or far away (smaller points) they are along dimension 1, which comes in and out of the page. (C) 3-D representation of the similarity of samples across all three dimensions. The size of points indicate distance from the viewer on the dimension-2 axis. In panels (A) and (B), points are shown with +/− standard error across the two dimensions shown on X and Y axes.

### *Wolbachia* colonization results in the differential expression of many cellular pathways

Differential expression analyses revealed 237 loci that were significantly differentially expressed due to *Wolbachia* colonization, regardless of time and virus infection (Supplementary Figure S2, Supplementary Table S2). Of these, 123 were upregulated and 114 were downregulated in the *Wolbachia*-colonized flies. We also detected significant differences is isoform usage due to *Wolbachia*: 8 of the differentially expressed genes (DEGs) also displayed differential isoform usage, and an additional 48 genes displayed differential isoform usage without any significant changes in the overall level of gene expression (Supplementary Tables S2 and S6). Changes in isoform usage included both changes to exon usage, and/or the transcribed regions of 3’ and 5’ UTRs. In total, 285 genes were either differentially expressed and/or displayed differential isoform usage.

We identified a core set of these 285 differentially expressed genes/isoforms that were predicted to interact with each other (Figure 2). Annotation of the core network revealed distinct processes and pathways that have perturbed gene expression patterns associated with *Wolbachia* colonization. These include stress responses, ubiquitin-related processes, metabolic functions, transcription and translation, RNA binding and processing, and recombination and cell cycle checkpoint.

**Figure 2.**
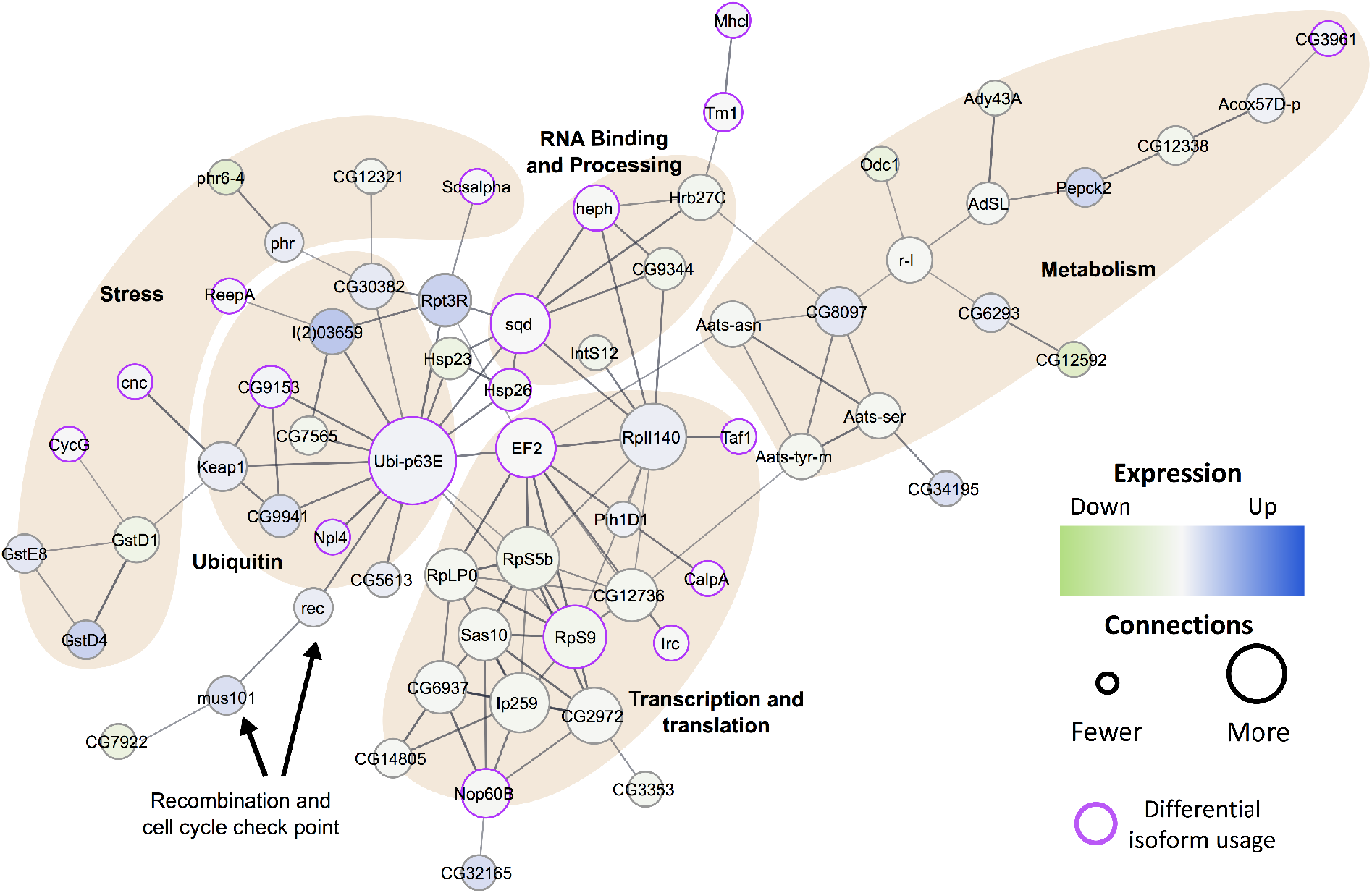
*Wolbachia*-responsive gene network comprises stress, ubiquitin, RNA binding and processing, transcription and translation, and metabolism pathways. STRING was used to identify the core network of interactions at a confidence threshold of 0.6 (relatively stringent). The size of a node corresponds to the number of connections to other nodes in the network. The color of a node corresponds to the level of expression relative to *Wolbachia*-free samples, where dark blue is up-regulation of gene expression, and light green is down-regulation of gene expression. Purple outlines indicate significant differences in transcript usage due to *Wolbachia* colonization. The weights of edges connecting the nodes indicate confidence of the interaction. Functional categories are indicated using shaded regions.

### Host response to virus infection varies depending on time and *Wolbachia* colonization

We identified 157 genes that were significantly differentially expressed due to virus infection (Supplemental Table S3). For 15 of these genes, time also had significant interactive effect on their level of expression, which is consistent with the MDS analyses (Supplemental Table S4). Virus infection resulted in significant differences in isoform usage for 38 genes, two of which were also significantly differentially expressed at the gene level (Supplemental Table S6). In total, 193 genes were differentially expressed and/or displayed differential isoform usage due to SINV. Again, we clustered genes with significant differences in expression due to virus, or virus*time based on their predicted interactions and identified a core network of genes (Figure 3). In contrast to the *Wolbachia* colonization core network, we find only two major functional categories represented in the SINV network: endoplasmic reticulum associated processes, and metabolic processes (mostly purine, sarcosine, and carbohydrate) (Figure 3).

**Figure 3.**
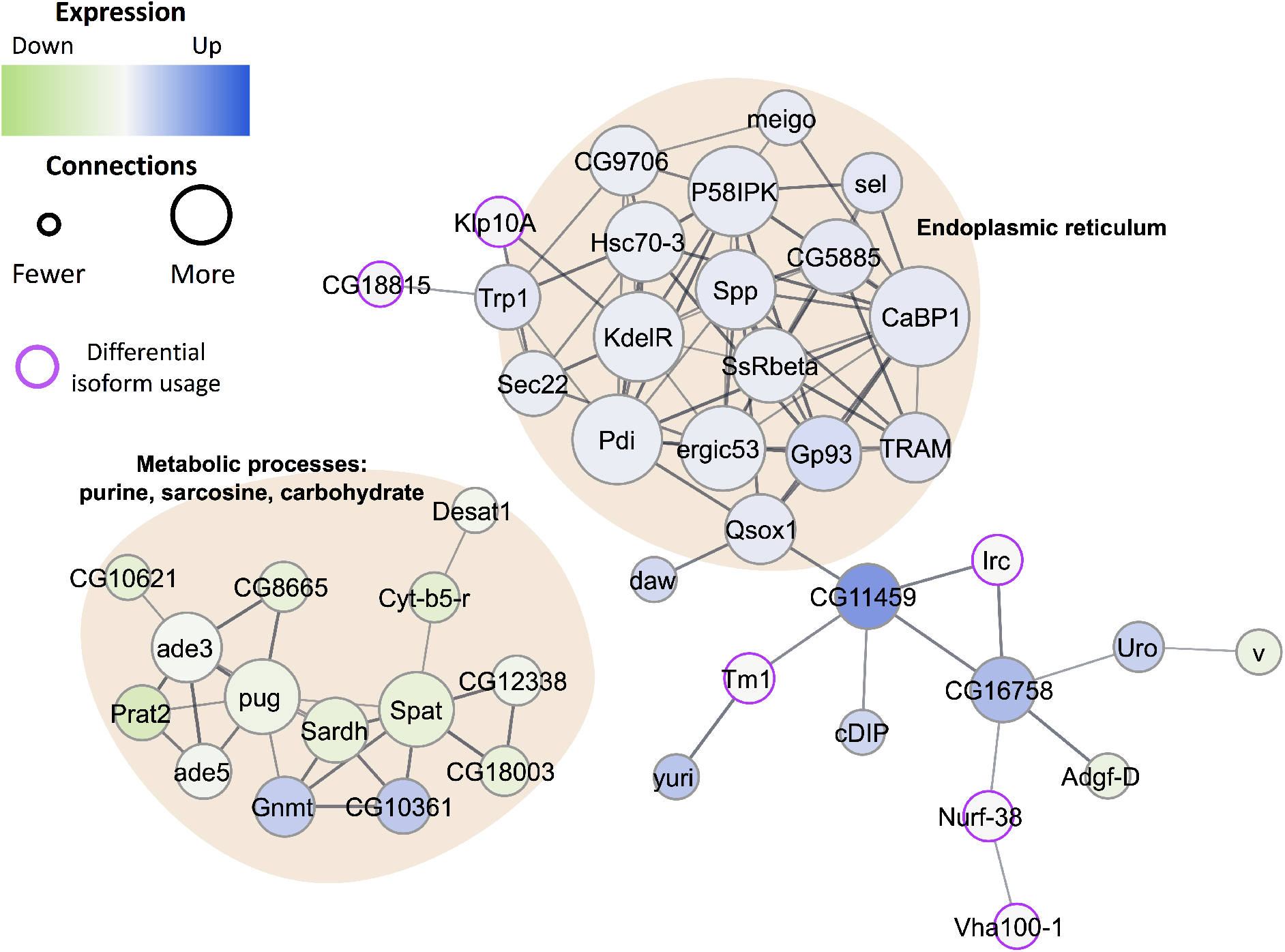
SINV-responsive core network comprises metabolic processes and endoplasmic reticulum pathways. STRING was used to identify the core network of interactions between proteins at a confidence threshold of 0.6 (relatively stringent). The size of a node corresponds to the number of connections to other nodes in the network. The color of a node corresponds to the level of expression relative to SINV-free samples, where dark blue is up-regulation of gene expression, and light green is down-regulation of gene expression. Purple outlines indicate significant differences in transcript usage due to SINV-infection. The weights of edges connecting the nodes indicate confidence of the interaction. Functional categories are indicated using shaded, annotated regions.

While there are limited genes that had significant changes in expression due to virus*time on a per-gene basis, it is clear that global expression patterns of all the virus-responsive DEGs vary across the duration of the experiment, which is consistent with the recovery patterns identified in the MDS plots (Figure 1). Additionally, while we did not identify any individual genes with altered expression due to the interaction of *Wolbachia* and virus, it is clear that on a global level, *Wolbachia*-free flies responded more dramatically to virus infection (Figure 4, Supplemental Figure S3). For DEGs that were upregulated upon virus infection, it was significantly more likely that any given upregulated DEG was more highly expressed in the *Wolbachia*-free flies than the *Wolbachia*-colonized flies (χ^2^ = 86.26, df = 2, p < 0.0001). It should be noted that these differences are subtle enough on a per-gene basis that they would not meet the criteria for an interactive effect of *Wolbachia* and virus, but across the set of upregulated DEGs we identified significant differences in the average log-fold change in gene expression between *Wolbachia*-free and *Wolbachia*-colonized flies. Upregulated DEGs were significantly more highly expressed due to the interaction of *Wolbachia* colonization and time post infection (ANOVA: F_1,650_ = 4.687, p = 0.0308). There were also significant effects of *Wolbachia* alone (ANOVA: F_1,650_ = 5.668, p = 0.0176), and time alone (ANOVA: F_1,650_ = 29.123, p < 0.0001). Indeed, 6 hpi, for *Wolbachia*-free flies, upregulated virus-responsive DEGs had an average of a 3.79-fold increase in expression relative to flies without virus, whereas *Wolbachia*-colonized flies on average experienced only a 2.57-fold increase in expression of the same DEGs (Figure 4B, Supplemental Figure S3A). This result suggests that *Wolbachia* colonization results in a muted host response to virus infection.

**Figure 4.**
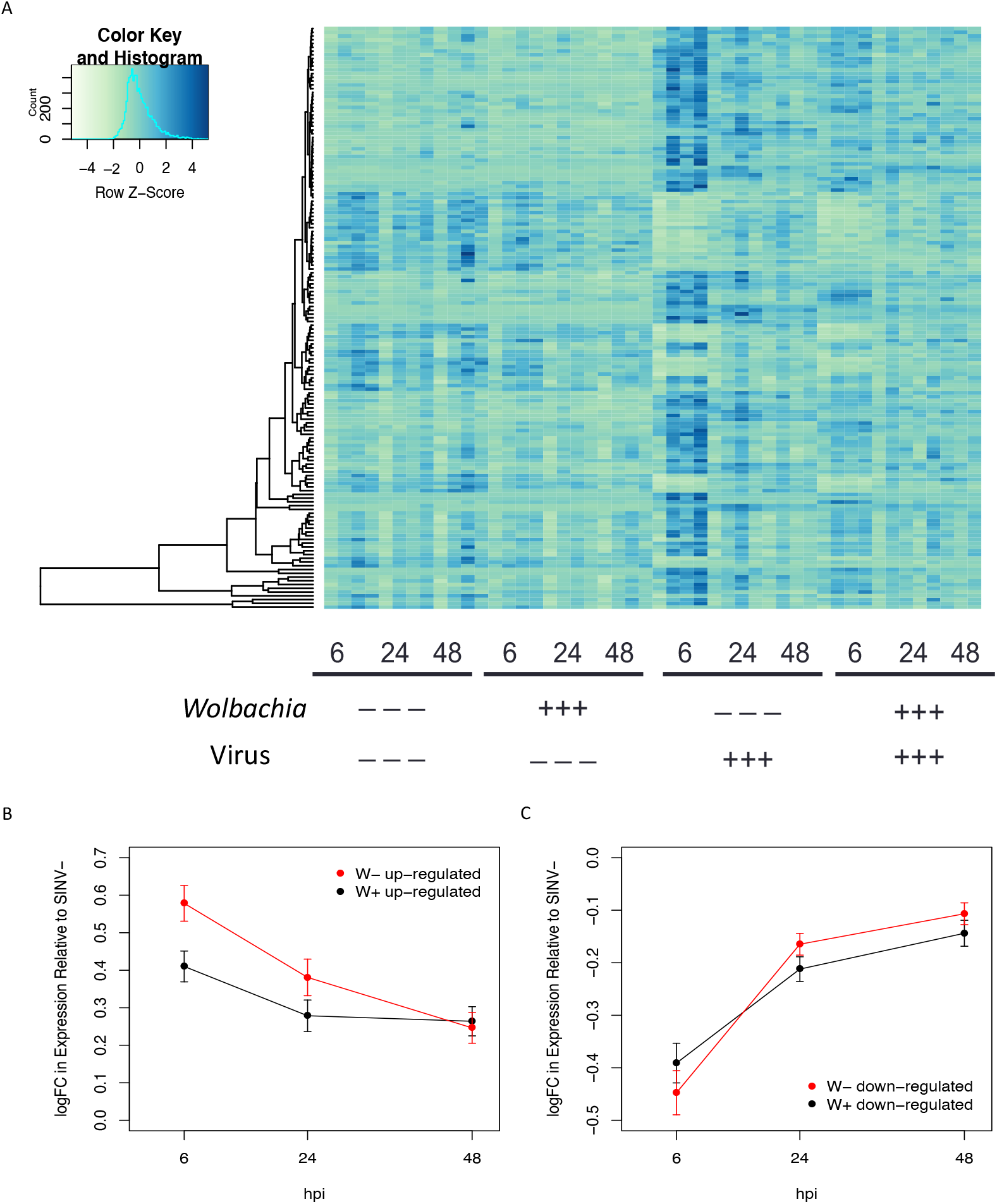
*Wolbachia* colonization alters the magnitude of response to virus infection. (A) Heatmap of the 157 genes significantly differentially expressed at the gene level, in response to SINV-infection at an FRD adjusted p-value of 0.05 and a fold change >2. *Wolbachia*-colonization, SINV-infection and timepoint are indicated under each set of samples, with biological replicates adjacent to each other. (B-C) Average log(FoldChange) in gene expression of virus-responsive genes for W+ (black) and W- (red) samples, relative to SINV-free flies. (B) Genes upregulated upon SINV infection. (C) Genes downregulated upon SINV-infection.

In contrast, the interaction between *Wolbachia*-colonization and time did not affect down-regulated, virus-responsive DEGs (ANOVA: F_1,272_ = 2.100, p = 0.1480). Additionally, *Wolbachia*-colonization alone had no significant effect on the change in expression of downregulated virus-responsive DEGs (ANOVA: F_1,272_ = 0.142, p = 0.7070). In other words, genes that were downregulated in response to virus infection did not show a significant effect based on *Wolbachia* colonization. Time post infection was the only factor that had a significant effect on the level of DEG expression (ANOVA: F_1,272_ = 88.345, p < 0.0001), as downregulated DEGs were most strongly downregulated at 6 hpi, and expression levels increased as flies recovered (Figure 4C, Figure S3B). However, we likely only see differences in the magnitude of response between *Wolbachia*-colonized and *Wolbachia*-free flies for upregulated DEGs and not downregulated genes due to decreased expression being bound by zero (or, no expression).

### *Wolbachia*-responsive and virus-responsive networks interact

While we did not identify any genes with expression levels that changed due to the interaction of *Wolbachia* and virus (either including or excluding the time factor), we identified 34 genes that responded interactively to *Wolbachia*virus* and/or *Wolbachia*\*virus*time at the level of isoform usage (Table 1, Supplemental Tables S6 and S7). These genes with interactive effects at the level of splicing were also significantly differentially expressed due to either *Wolbachia* or virus alone. These 34 differentially spliced genes include a range of predicted functions including transcription and translation (*eEF2, MED26*, and *da*), cytoskeletal organization (*sickie, CAP, Eb1, hts*, and *Klp10A)*, nucleotide metabolic processes (*Pde11)*, and immune and stress responses (*Irc* and *cert)*, amongst others (Table 1, Supplemental Tables S6 and S7).

**Table 1.** Genes with isoform usage patterns that were significantly affected by the interaction of virus and *Wolbachia*.

| FlyBase ID | Gene | Notes | Abbreviated GO Annotations* |
| --- | --- | --- | --- |
| FBgn0000559 | eEF2 |  | translational elongation |
| FBgn0001225 | Hsp26 |  | stress responses |
| FBgn0004509 | Fur1 | *Wolbachia* suppressor (Grobler et al., 2018) | neurotransmitter and protein processing |
| FBgn0013765 | cnn | *Wolbachia* enhancer (Grobler et al., 2018) | mitotic spindle organization |
| FBgn0016687 | Nurf-38 |  | chromatin remodeling, signaling |
| FBgn0020370 | TppII |  | proteolysis |
| FBgn0023522 | CG11596 |  | carnosine metabolic process |
| FBgn0026415 | Idgf4 |  | chitin metabolism |
| FBgn0027066 | Eb1 |  | spindle organization and elongation; sensory development and locomotion |
| FBgn0027569 | cert |  | sphingolipid metabolism and transport |
| FBgn0030087 | CG7766 |  | glycogen metabolism and protein phosphorylation |
| FBgn0030268 | Klp10A |  | spindle organization |
| FBgn0030503 | Tango2 |  | Golgi organization and protein secretion |
| FBgn0030504 | CG2691 |  | N/A |
| FBgn0032906 | RPA2 |  | DNA repair |
| FBgn0033504 | CAP |  | sensory perception |
| FBgn0034075 | Asph | *Wolbachia* enhancer (Grobler et al., 2018) | peptidyl-aspartic acid hydroxylation |
| FBgn0036932 | CG14184 |  | endomembrane system transport and localization |
| FBgn0037810 | sle |  | nucleolus organization |
| FBgn0037944 | CG6923 |  | ubiquitin-dependent protein catabolic process |
| FBgn0038465 | Irc |  | oxidation-reduction process |
| FBgn0038470 | CG18213 |  | N/A |
| FBgn0038535 | alt | *Wolbachia* enhancer (Grobler et al., 2018) | N/A |
| FBgn0039350 | jigr1 | *Wolbachia* enhancer (Grobler et al., 2018) | regulation of gene expression |
| FBgn0039466 | CG5521 |  | regulation of GTPase activity |
| FBgn0039923 | MED26 |  | regulation of gene expression |
| FBgn0042138 | CG18815 |  | protein depalmitoylation |
| FBgn0043799 | CG31381 |  | tRNA modification |
| FBgn0052264 | CG32264 |  | actin cytoskeleton reorganization |
| FBgn0053193 | sav |  | signaling and growth regulation |
| FBgn0085370 | Pde11 |  | signal transduction |
| FBgn0263391 | hts |  | meiotic spindle organization, actin organization |
| FBgn0263873 | sick | pro-vial (Panda et al., 2013) | actin organization, nervous system development, response to bacterium |
| FBgn0267821 | da |  | development and cell differentiation |

\* Full annotations in Supplemental Table S7

Next, we clustered all infection responsive genes (either at the level of gene expression and/or isoform usage) to determine how interconnected the *Wolbachia*- and virus-responsive genes sets are. Each gene was classified as either “*Wolbachia*-responsive”, “virus-responsive”, “interaction-responsive” (the 34 for genes mentioned above), or those affected by both *Wolbachia* and SINV, but non-interactively (for example, differentially expressed due to *Wolbachia* colonization, and differential isoform usage due to SINV infection). We identified one core network that includes genes across all responses, with numerous connections between *Wolbachia*-responsive, virus-responsive, and interactive response genes (Figure 5). This clustering revealed that metabolic processes are the most interconnected between the different responses, particularly *de novo* nucleotide synthesis. Indeed, we identified numerous GO Processes that were significantly enriched in the joint network, all of which were metabolic in nature (Supplemental Table S8). Enrichments included amino acid metabolic processes, purine biosynthesis, and other small molecule metabolic processes.

**Figure 5.**
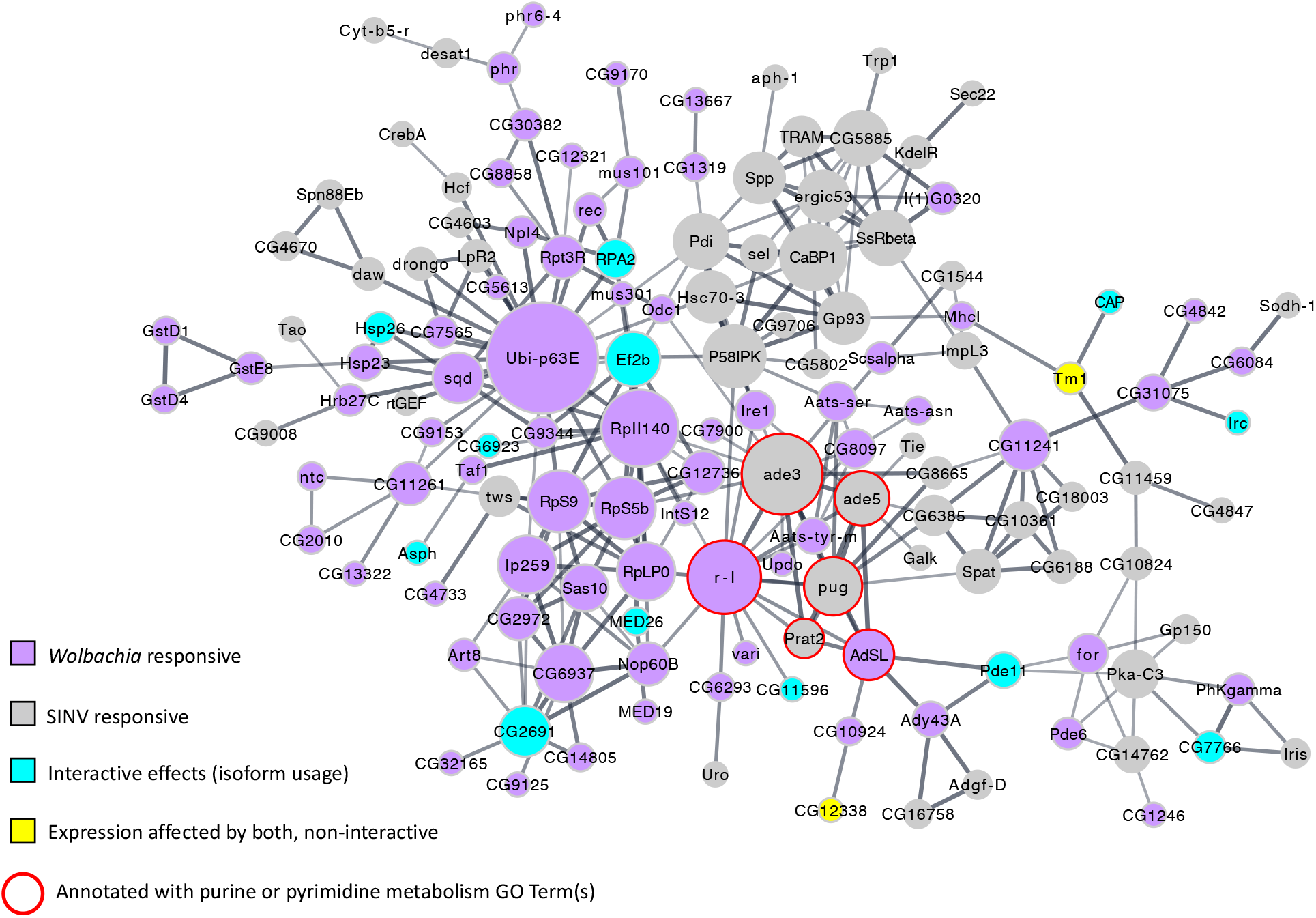
Virus and *Wolbachia* responsive genes are enriched for metabolic processes and interconnected around nucleotide metabolism. All infection-responsive genes were clustered to look for connectivity of the *Wolbachia*-responsive and virus-responsive gene sets. Node size indicates the number of connections to other nodes in the network. Purple nodes are *Wolbachia*-responsive, grey nodes are virus-responsive. Interactive effects on expression are indicated by blue nodes. The only significant interactive effects were differential transcript usage, so all blue nodes are differentially spliced. Yellow nodes indicate genes were both SINV and *Wolbachia* had significant effects on expression, but the effect was not interactive.

### Nucleotide metabolism and *Wolbachia* colonization have interactive effects on virus replication

Given the interconnectedness of the infection responsive networks around nucleotide metabolic processes (Figure 5), we used fly genetics to determine if these changes in gene expression were pro- or antiviral. First, used the RNA-Seq data to determine how *Wolbachia* colonization and virus infection affected expression of the entire *de novo* purine and pyrimidine synthesis pathways (Figure 6A-C). These pathways are directly connected (an intermediate product of purine synthesis is required for a step of pyrimidine synthesis; Figure 6A), and the expression of many genes encoding for enzymes involved in the pathway are significantly altered by *Wolbachia* or virus (Figure 5). In general, the purine synthesis pathway is strongly downregulated due to virus (Figure 6B), and the pyrimidine synthesis pathway is strongly downregulated due to *Wolbachia* (including upregulation of a suppressor, *su(r)*)(Figure 6C). Interestingly, there are a few genes that differentially respond to *Wolbachia* and virus, such as *prat2. prat2* is a gene involved in the *de novo* synthesis of purine nucleotides (Ji and Clark, 2006; Malmanche et al., 2003), and one of the most strongly downregulated in the virus-responsive gene set, expressed at <0.01% of the level of expression in PBS-injected flies (Figure 6B). While *prat2* did not meet the threshold for statistical significance in the *Wolbachia*-responsive RNA-Seq analysis, *prat2* was upregulated in *Wolbachia*-colonized flies 1.7-fold (Figure 6B).

**Figure 6.**
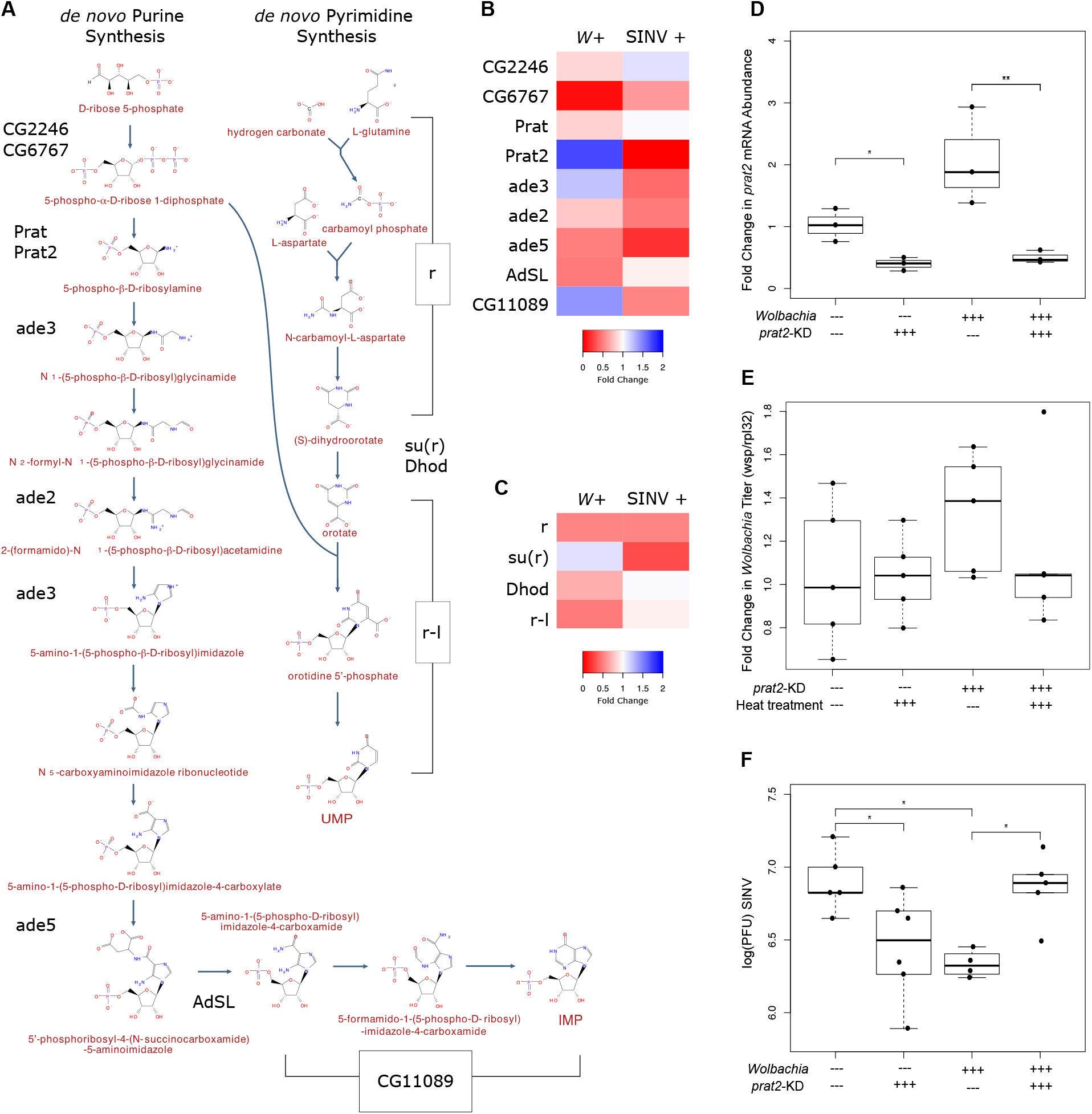
*Wolbachia* and nucleotide metabolism genes have interactive effects on virus replication. **(A)** *de* novo biosynthesis of purines (inosine monophosphate; IMP) and pyrimidines (uridine monophosphate; UMP) in *Drosophila melanogaster*. Genes encoding for enzymes are in black. **(B)** Change in gene expression of *de novo* purine synthesis genes due to *Wolbachia* colonization (*W*+) and virus infection (SINV+). **(C)** Change in gene expression of *de novo* pyrimidine synthesis genes due to *Wolbachia* colonization (*W*+) and virus infection (SINV+). **(D)** *prat2* mRNA levels were quantified in flies with or without *Wolbachia (W+/W-)*, that did or did not contain a *prat2* silencing shirt hairpin RNA (TRiP-*prat2* or sibling, respectively), using qRT-PCR, relative to the expression of rpl32. Both *Wolbachia* and the presence of the shRNA resulted in significant differences in *prat2* expression (ANOVA: *Wolbachia:* F_1,8_ = 7.659, p = 0.0244; TRiP-*prat2*: F_1,8_ = 48.697, p = 0.0001). In sibling controls (no knockdown) *prat2* expression in *Wolbachia*-colonized flies was on average 2.06-fold higher than *Wolbachia*-free flies. In both *Wolbachia*-colonized and *Wolbachia*-free flies, *prat2* knockdown was effective, resulting in *prat2* mRNA levels being reduced to 24.3% and 39.6% of the sibling controls with the same *Wolbachia* colonization status. There was no significant difference in *prat2* mRNA levels between *Wolbachia*-colonized and *Wolbachia*-free flies with the shRNA (Tukey’s, p = 0.7271). **(E)** Flies with *Wolbachia* that did or did not contain the *prat2* targeting shRNA were either heat shocked (10 minutes 37C), or not, to determine if heat and/or presence of the shRNA had an effect on *Wolbachia* titer, which might affect downstream pathogen blocking efficiency. Neither heat nor the presence of the shRNA resulted in significant differences in *Wolbachia* titer (ANOVA: TRiP-*prat2*: F_1,16_ = 1.985, p = 0.178; heat: F_1,16_ = 0.451, p = 0.512). **(F)** Flies with and without *Wolbachia*, and with or without knockdown of *prat2* were injected with SINV to assess the effect of *Wolbachia* and *prat2* on virus replication. Viral titers from whole flies were assessed with standard plaque assays on BHK-21 cells. *Wolbachia* colonization and *prat2* knockdown had a significant interactive effect on SINV titers (ANOVA: F_1,16_ = 17.633, p = 0.0007). Sibling controls (no shRNA) with and without *Wolbachia* recapitulated the pathogen blocking phenotype, with SINV titers significantly reduced, by approximately half a log, in the *Wolbachia*-colonized flies (Tukey’s, p=0.0217). When *prat2* titers were knocked down, there was a *Wolbachia*-colonization-dependent effect on SINV, with knockdown being significantly pro-viral in the presence of *Wolbachia* (Tukey’s, p=0.0350), and antiviral in the absence of *Wolbachia* (Tukey’s, p= 0.0469).

We used transgenic RNAi fly lines to knockdown *prat2* gene expression and assess the effect on SINV replication. Knockdown was achieved using a *prat2*-targeting short-hairpin RNA (shRNA), with expression induced using heat-shock conditions (Hsp.70-GAL4 driving UAS-anti-prat2). Sibling controls without the *prat2*-targeting shRNA recapitulated the increase in *prat2* expression seen in *Wolbachia*-colonized flies (here, ~2-fold increase and statistically significant (ANOVA: F_1,8_ = 7.659, p = 0.0244)), similar to what was observed in the RNA-Seq dataset (1.7-fold increase). In both *Wolbachia*-colonized and *Wolbachia*-free flies, *prat2* knockdown was effective, resulting in *prat2* mRNA levels being reduced to 24.3% and 39.6% of the sibling controls, respectively (Figure 6D). There was no significant difference in *prat2* mRNA levels between *Wolbachia*-colonized and *Wolbachia*-free flies with the shRNA (Tukey’s, p = 0.7271). Neither heat shock nor knockdown had a significant effect on *Wolbachia* titer (Figure 6E; ANOVA: TRiP-*prat2*: F_1,16_ = 1.985, p = 0.178; heat: F_1,16_ = 0.451, p = 0.512), which is known to affect the efficiency of pathogen blocking (Osborne et al., 2012; Osborne et al., 2009). 24-hours post knockdown, flies were injected with SINV to determine the effect of *prat2* expression and *Wolbachia* on virus titer. *Wolbachia* colonization and *prat2* knockdown had a significant interactive effect on SINV titers (Figure 6F: ANOVA: F_1,16_ = 17.633, p = 0.0007). Sibling controls (no shRNA) with and without *Wolbachia* recapitulated the pathogen blocking phenotype, with SINV titers significantly reduced, by approximately half a log, in the *Wolbachia*-colonized flies (Tukey’s, p=0.0217), typical of what has previously been seen in this system (Bhattacharya et al., 2017). When *prat2* expression was knocked down, there was a *Wolbachia*-colonization-dependent effect on SINV replication, with knockdown being significantly proviral in the presence of *Wolbachia* (Tukey’s, p=0.0350), and antiviral in the absence of *Wolbachia* (Tukey’s, p= 0.0469), indicating that nucleotide metabolic processes are likely a point of interaction between host, *Wolbachia*, and virus in this system.

## DISCUSSION

*Wolbachia* colonization is well known for altering numerous physiological processes in its hosts. In many *Wolbachia*-host associations this appears to have an effect on secondary infections, mainly with RNA viruses. Given that many different processes have been implicated in resistance to RNA viruses resulting from *Wolbachia*-colonization (Lindsey et al., 2018a), and it has been hypothesized that the preexisting state of cells with *Wolbachia* is responsible for reduced virus replication (Lindsey et al., 2018a), we used a model system to better explore the *Wolbachia*-host relationship. We identified changes in both gene expression and isoform usage due to *Wolbachia* colonization in whole flies and identified key processes that are perturbed as a result of *Wolbachia*. This deeper look into the association allowed us to more efficiently overlay the changes that occur due to virus, and identify areas of overlapping effects, regardless of whether or not they were combinatorial.

One of the major findings across our analyses is the significant amount of differential isoform usage due to *Wolbachia*, virus, and the combination of the two. The first evidence of *Wolbachia* having effects on host splicing and/or isoform usage was recently reported in a parasitoid wasp (Wu et al., 2020), and splicing is becoming increasingly appreciated as an important component of host-microbe interactions (Chauhan et al., 2019; Rotival et al., 2019). Whether or not *Wolbachia* directly modulates splicing via secreted factors or splicing is a host response to infection is yet to be determined, but there are likely to be many downstream effects due to changes in isoform usage and stoichiometry.

We find that *Wolbachia* colonization affects the expression of many different biological processes, including: (A) stress responses, (B) ubiquitination, (C) transcription and translation, (D) RNA binding and processing, (E) metabolism, and (F) cell cycle checkpoint and recombination. Many of these have been previously explored in host-*Wolbachia* relationships in more targeted studies (e.x., ROS and stress (Brennan et al., 2012; Pan et al., 2012; Wong et al., 2015), translation (Grobler et al., 2018)), and/or agree with previously identified effects that *Wolbachia* has on the host (e.x., the CI genes encode a deubiquitylase (Beckmann et al., 2017; Lindsey et al., 2018b)). The effects of *Wolbachia* on host metabolism are arguably underexplored (Newton and Rice, 2020), which is surprising given that *Wolbachia* must acquire all nutrients from the host, encodes for a select number of its own metabolic pathways, and encodes for a variety of transporters that would allow for *Wolbachia* to import specific metabolites (e.x., amino acids) (Wu et al., 2004).

We next identified the changes in gene expression and isoform usage due to the presence of virus. The virus-responsive network contained fewer cellular processes than did the *Wolbachia*-responsive network: the response to virus mainly affected the expression of endomembrane system associated genes, and metabolic pathways. It is notable that these metabolic pathways are largely distinct from the *Wolbachia*-responsive metabolic pathways at the gene level, though there is likely the potential for interaction at the level of metabolites and flux in the cell.

Proteins and metabolites directly involved in blocking need not be differentially expressed or differentially abundant to result in the decreased replication of virus. *Wolbachia’s* restructuring of the intracellular space could lead to changes in localization, modification, or the availability of co-factors and substrates that may be critical for the expression of an antiviral effect (Lindsey et al., 2018a). Furthermore, it is important to note that many of the processes previously identified as being involved in the blocking phenotype are (A) not mutually exclusive, (B) have the potential to act at different points in the virus life cycle, and (C) may be up or downstream from each other in a network of cellular changes that ultimately affect virus replication.

While it is likely that there are key differences in the mechanism(s) of pathogen blocking between different *Wolbachia*-host-virus associations, it does not exclude the possibility that there are similar up-stream events (ex., *Wolbachia* using host amino acids) that result in dissimilar downstream events that are dependent upon both (A) the host, and (B) the combination of other *Wolbachia*-induced changes in physiology (ex., a host immune response to *Wolbachia’s* presence, which is more common in nonnative *Wolbachia*-host associations (Lindsey et al., 2018a)). In many previously published studies, a pathway has been implicated in pathogen blocking but it was not determined how the change in host physiology occurred, and whether or not that effect was directly or indirectly responsible for blocking. For example, what results in changes to expression of the Toll pathway? AMPs are differentially expressed, but do they have a direct effect on virus replication? Or, do the AMPs act as signaling molecules that in turn alter other host processes? Changes in lipid abundance have been associated with the antiviral effect (Caragata et al., 2013; Geoghegan et al., 2017; Molloy et al., 2016), but it is unclear if this perturbation in lipids results in other changes to host gene expression or cellular structure, or if the virus particles themselves are unable to properly form their membrane-associated replication factories or envelopes during assembly.

Here, we identified the expression of metabolic processes as significantly altered due to both *Wolbachia* and virus. This agrees with previously published studies (Newton and Rice, 2020) and what we know about *Wolbachia* and virus biology. *Wolbachia* encode a suite of amino-acid importers, which likely results in altered amino acid pools in the host (Wu et al., 2004). Amino acids are not only critical for protein synthesis of course, but also serve as precursors for many metabolic processes, including the *de novo* synthesis of purine and pyrimidine nucleotides. Here, we used fly genetics to explore the effect of *de novo* purine synthesis gene expression on the *Wolbachia*-virus-host relationship. Not only did *prat2* gene expression have an effect on viral titers, it was dependent on the presence of *Wolbachia*, which further highlights the complexity of the system and implicates multiple processes in the pathogen blocking phenotype. In *Wolbachia*-colonized flies, where *prat2* is upregulated, *prat2* knockdown was pro-viral which supports the idea of the pre-existing state of *Wolbachia*-colonized flies being antiviral. It is unclear what the downstream effects of altered *prat2* expression are, and how they may be different between flies with and without *Wolbachia*. For example, knockdown of the *de novo* purine synthesis pathway may result in increased expression of the purine salvage pathway. The biochemical reactions for these different pathways have different byproducts and intermediates which may have an effect on other cellular processes and virus. These downstream consequences of *prat2* knockdown may be the reason we see interactive effects of *Wolbachia* presence and *prat2* expression on virus titer.

The finding that nucleotide metabolism is a source of interaction between *Wolbachia* and virus is particularly interesting given that many currently marketed antiviral drugs are known to interfere with nucleotide metabolic processes, often in the same pathways that we identify here as being perturbed due to *Wolbachia* and/or virus. For example, Ribavirin and other compounds confer broad spectrum antiviral activity by inhibition of IMP dehydrogenase, an enzyme involved in purine metabolic processes (Leyssen et al., 2005; Markland et al., 2000). The antiviral activity of another compound, Favipiravir, is reduced in the presence of excess purines (Furuta et al., 2002). A more recently identified broad spectrum antiviral was shown to interfere with pyrimidine metabolism via dihydroorotate dehydrogenase (Hoffmann et al., 2011b) (*dhod* in *Drosophila)*, which was significantly differentially expressed due to *Wolbachia* colonization in our study. Similarly, an excess of pyrimidines rescues virus replication in the presence of this antiviral compound. A separate group of antivirals, brequinar, leflunomide, and derivatives, are also known to interfere with dihydroorotate dehydrogenase and pyrimidine pools, which is responsible for the broad spectrum antiviral effect (Greene et al., 1995; McLean et al., 2001).

Additional studies are needed to determine the effect of *Wolbachia* and virus on the nucleotide pools of host cells, but it is plausible that this is a major source of conflict between these two intracellular inhabitants. Indeed, metabolomic analyses will likely provide a wealth of information that will help us connect transcriptomic changes to downstream events in the physiology of the host that eventually result in a pathogen blocking phenotype.

## ACKNOWLEDGEMENTS

This work was supported by the National Institute of Allergy and Infectious Disease (R21 AI121849 to ILGN and RWH, and R01AI144430 to ILGN). Stocks obtained from the Bloomington *Drosophila* Stock Center (NIH P40OD018537) were used in this study. Thank you to MaryAnn Martin, Audrey Parish, Delaney Miller, Lindsay Nevalainen, and Danny Rice for comments on an earlier draft of the manuscript.

## AUTHOR CONTRIBUTIONS

Conceptualization, I.L.G.N. and R.W.H.; Methodology, A.R.I.L. and T.B.; Investigation, A.R.I.L. and T.B.; Formal Analysis, A.R.I.L.; Visualization, A.R.I.L.; Writing – Original Draft, A.R.I.L.; Writing – Reviewing & Editing, A.R.I.L., T.B., I.L.G.N., and R.W.H.; Funding Acquisition, I.L.G.N. and R.W.H.; Resources, I.L.G.N. and R.W.H.; Supervision, A.R.I.L., I.L.G.N., and R.W.H.

## METHODS

### *Drosophila* husbandry

A previously described line of *Drosophila melanogaster*, stock 6326 from the Bloomington *Drosophila* Stock Center (http://flystocks.bio.indiana.edu/), a *w*^1118^ background infected with *Wolbachia* strain *w*Mel2, and its *Wolbachia*-cleared counterpart, were used in transcriptomic experiments (Bhattacharya et al., 2017). *Wolbachia* colonization status was confirmed using specific primers that target the *Wolbachia*-specific *wsp* locus (Baldo et al., 2006). Fly stocks were maintained on standard cornmeal-agar medium at 25 °C on a 24-hour light: dark cycle under density-controlled conditions.

### Cell culture and virus preparation

BHK-21 cells (American Type Culture Collection) were grown at 37 °C under 5% CO2 in MEM (CellGro) supplemented with 1% L-Gln, 1% Antibiotic-Antimycotic (Gibco), 1% non-essential amino acids and 10% heat inactivated Fetal Bovine Serum (FBS)(Corning™). SINV (strain TE3’2J-GFP (Huang et al., 2013)) was prepared by transfecting Baby Hamster Kidney Fibroblasts (BHK-21) cells with 1μg of *in vitro* transcribed viral RNA with Lipofectamine LTX (Sigma Aldrich) to generate a P0 virus stock, which was then used to infect new BHK-21 cells to generate P1 virus (Huang et al., 2013). The supernatant containing P1 virus was collected, purified by centrifugation over a 27% (w/v) sucrose cushion in 1XHNE buffer (20mM HEPES, 0.15M NaCl, 0.1mM EDTA), resuspended in 1XHNE, and viral titers were determined by standard plaque assays on BHK-21 cells as done previously (Huang et al., 2013).

### *Drosophila* injections

To determine the effect of *Wolbachia* and virus infection on fly gene expression, and the effect of virus on *Wolbachia* gene expression, we established *in vivo* systemic viral infections in adult *Drosophila*, using a block design with a time series. Flies with or without *Wolbachia* (W+/W-), were injected with either virus or saline (SINV+/SINV-), and collected at 6, 24, and 48 hours post-injection (hpi). For each unique condition of W-SINV-time, we generated four biological replicates (A-D), with each replicate consisting of a pool of five virgin females. Specific conditions for generating the fly infection conditions are as follows: five-day-old virgin female *Drosophila* were anesthetized with CO2 and injected with either: (a) 50 nl sterile phosphate buffered saline (PBS), or (b) 50 nl of freshly grown SINV (10^10^ PFU/mL) using a nanoinjector (Drummond Scientific). Pools of five flies (representing a single biological replicate) were injected in a randomized order across a five-hour time period, and capillary needles were changed between fly types (*Wolbachia*-colonized or not) and injection type (PBS or SINV) to avoid cross-contamination. Exact time of injection was recorded, and the pool of five females was placed in a vial containing standard cornmeal-agar medium supplemented with antibiotic-antimycotic (Corning) and a fresh Kimwipe. Subsequently. 6, 24, or 48 hpi flies were flash frozen in liquid nitrogen and stored at −80 °C until further processing.

### RNA extractions, library preparation, and sequencing

RNA was extracted from pools of flash frozen flies using TRIzol™ Reagent (Invitrogen) following bead-beating, and according to manufacturer’s instructions. rRNAs and other uncapped RNA species were depleted from RNA samples using the Terminator™ 5’-Phosphate-Dependent Exonuclease (Lucigen). Following a standard phenol-chloroform-isoamyl precipitation, cDNA libraries were prepared with the NEBNext® Ultra™ II Directional RNA Library Prep Kit (New England Biolabs) following manufacturer’s recommendations, including a seven minute fragmentation time, 10 cycles of PCR amplification, and use of a specific barcode from the NEBNext® Multiplex Oligos for Illumina® Index Primers Set 1 or 2 (New England Biolabs). Quality and quantity of total RNA, depleted RNA, and final libraries was assessed using a TapeStation 2200 (Agilent). Libraries were pooled in groups of 16 such that biological replicates of *Wolbachia* colonization status, SINV infection status, and time were split as evenly as possible across three runs on an Illumina® NextSeq to generate 75 base pair single-ended ended reads. An average of 32.5 million reads were generated for each library. Further details and mapping statistics can be found in Supplemental Table S1.

### Transcriptomic analyses

Following demultiplexing, reads were mapped to extracted reference transcripts of either the *Drosophila melanogaster* reference genome (release 6.16) (dos Santos et al., 2014) or the *w*Mel strain *Wolbachia* genome (GenBank accession NC_002978.6 (Wu et al., 2004)) using the RSEM v. 1.3.0 (Li and Dewey, 2011) programs ‘rsem-prepare-reference’ and ‘rsem-calculate-expression’, employing the default Bowtie aligner (Langmead and Salzberg, 2012). Transcript abundance was summarized and imported to R v. 3.3.1 ‘Bug in Your Hair’ (R Core Team, 2014) with tximport v. 1.2.0 (Soneson et al., 2015) for use in downstream analyses. Differential gene expression and splicing was assessed with EdgeR v. 3.16.5 (McCarthy et al., 2012; Robinson et al., 2010), employing a TMM normalization, dispersion calculation, and a multivariate generalized linear model (*‘~Wolbachia* * SINV * time’ for *Drosophila* expression, or ‘~SINV * time’ for *Wolbachia* expression), with quasi-likelihood F-tests (function ‘glmQLFit’). Splicing was assessed with the ‘diffSpliceDGE’ function using the ‘Simes’ method. Genes that were significantly up- or down-regulated were defined as those with a false discovery rate (FDR) q-value of <0.05. To check for SINV reads, libraries were mapped to the SINV TE3’2J-GFP (Huang et al., 2013) reference genome with BWA v.0.7.12 (Li, 2013) and mapping statistics were assessed with samtools v.1.5 (Li et al., 2009). Raw reads will be made publicly available on NCBI GenBank (submission and accession number generation in process). Reviewers can request access to raw reads at this time.

### Statistics, data visualization, and network analysis

Statistical analyses and plotting were carried out in R v. 3.3.1 ‘Bug in Your Hair’ (R Core Team, 2014). Three-D plots were generated with the R package ‘plot3D’ (Soetaert, 2013), implementing the ‘scatter3D’ function. Protein-protein interaction networks were constructed with STRING v.1.4.2 (Doncheva et al., 2018), implemented in Cytoscape v.3.6.0 (Shannon et al., 2003). The confidence threshold for all networks was set to 0.600, considered stringent, so as to limit the complexity of the networks and identify the strongest interactions. Nucleotide biosynthesis pathway information was downloaded from BioCyc (Caspi et al., 2016).

### Nucleotide metabolism fly mutants

Given the altered expression of genes related to nucleotide metabolism in *Wolbachia*-host and SINV-host relationships (see Results), we chose to study key fly pathways to further define these relationships. Stocks were reared and screened for *Wolbachia* using the same protocols detailed above (see *“Drosophila* husbandry”). We used a fly stock which carries a UAS-*prat2*-specific short hairpin silencing trigger (BDSC stock #51492, RNAi TRiP line: y^1^sc*v^1^sev^21^; P{y^+t7.7^ v^+t1.8^=TRiP.HMC03244}attP2). This stock is *Wolbachia*-colonized. To generate balanced heterozygous offspring, we crossed virgin females homozygous for the hairpin to males with a 3^rd^ chromosome balancer (BDSC stock #6663: w^1118^; Dr^Mio^/TM3, P{w^+mC^=GAL4-twi.G}2.3, P{UAS-2xEGFP} AH2.3, Sb^1^Ser^1^). To knock down the expression of *prat2*, the *Wolbachia*-colonized, balanced flies carrying the UAS-anti-*prat2* insert were crossed to a *Wolbachia* free, homozygous, inducible Hsp70:Gal4-driver line (BDSC stock #2077: w*; P{w^+mC^=GAL4-Hsp70.PB}2). Crosses were performed in both directions to generate *Wolbachia*-free and *Wolbachia*-colonized offspring. Virgin females with and without *Wolbachia*, that contained either: (1) both the Gal4-driver and the UAS-anti-*prat2*, or (2) sibling controls with the driver and the TM3 balancer were collected and aged four days. At four days old, all flies were heat shocked at 37°C for 10 minutes to induce Gal4 expression. 24-hours post heat shock, flies were either injected with virus or flash frozen to asses knockdown of *prat2*, or *Wolbachia* titer. For virus infections, flies were injected intrathoracically with SINV following the same injection and recovery protocol as were used for setting up the transcriptomics experiment. 48 hpi, flies were harvested and viral titers from single flies were assessed with standard plaque assays on BHK-21 cells using technical duplicates of each fly (Bhattacharya et al., 2017).

### Real-time quantitative RT-PCR analyses of *prat2* expression

Single flies were homogenized in TRIzol™ Reagent (Invitrogen), and RNA was extracted and DNAse treated according to manufacturer’s instructions. *prat2* expression was assessed with the SensiFAST™ SYBR® Hi-ROX One-Step Kit (Bioline) according to manufacturer’s recommendations with specific primers PP25361 (Forward: 5’-GGGAATAGGACACACCCGGTA-3’; Reverse: 5’-GCAGTTCACTAGCTCACCATT-3’)(Perkins et al., 2015), and normalized to expression of Rpl32 (Forward: 5’-CCGCTTCAAGGGACAGTATC-3’; Reverse: 5’-CAATCTCCTTGCGCTTCTTG-3’ (Newton et al., 2015)) using the Livak method (Livak and Schmittgen, 2001). All samples were run in technical duplicate alongside a standard curve and negative controls on an Applied Bioscience StepOnePlus qRT-PCR machine (Life Technologies).

### Real-time quantitative PCR analyses of *Wolbachia* titer

DNA was extracted from single flies using Qiagen DNeasy Blood and Tissue Kit (Qiagen), according to manufacturer’s instructions. *Wolbachia* titer was determined by amplification of the single-copy *Wolbachia* gene, *wsp*, and normalized to abundance of the host gene *rpl32* according to previously established protocols (Newton et al., 2015) using PowerUp™ SYBR™ GreenMaster Mix (ThermoFisher). All samples were run in technical duplicate alongside a standard curve and negative controls on an Applied Bioscience StepOnePlus qRT-PCR machine (Life Technologies).

## SUPPLEMENTAL INFORMATION

**Supplemental Tables (Tabs in Excel File)**

**Supplemental Table S1.** Sequencing library statistics.

**Supplemental Table S2.** Transcriptomic response to *Wolbachia*. DEGs are those with corrected P values < 0.05.

**Supplemental Table S3.** Transcriptomic response to SINV. DEGs are those with corrected P values < 0.05.

**Supplemental Table S4.** Transcriptomic response to SINV*TIME. DEGs are those with corrected P values < 0.05.

**Supplemental Table S5.** All normalized counts for all genes across all samples.

**Supplemental Table S6.** Genes with significant differences in isoform usage according to explanatory variable coefficients.

**Supplemental Table S7.** Genes with significant differences in isoform usage due to *Wolbachia*\*SINV and/or *Wolbachia*\*SINV*TIME along with functional annotations and notes to references in previously published literature.

**Supplemental Table S8.** Enriched terms in the joint STRING network.

**Supplemental Figures (PDF)**

**Figure S1.** MDS plot of *Wolbachia* gene expression.

**Figure S2.** Transcriptomic response to *Wolbachia* colonization.

**Figure S3.** *Wolbachia* colonization results in a muted response to virus infection.

