## Supplemental Figures S1-3 for "*Wolbachia* and virus alter the host transcriptome at the interface of nucleotide metabolism pathways"

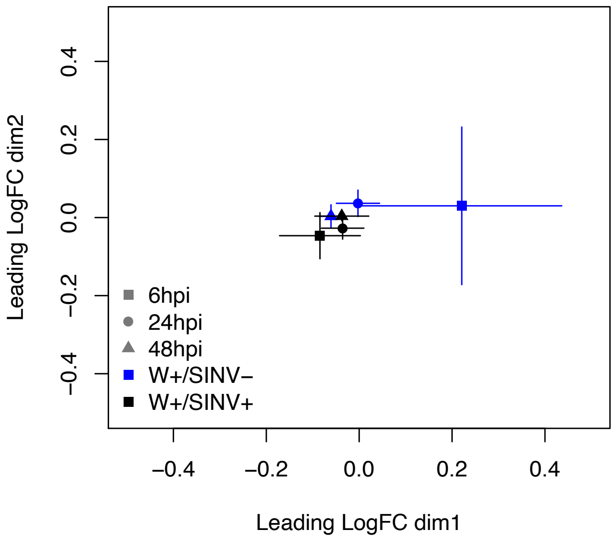


**Figure S1. MDS plot of *Wolbachia* gene expression.** MDS plot showing similarity of total gene expression across samples. Within each SINV-timepoint combination, biological replicates were averaged to show their center of gravity +/- standard error across dimension-1 and -2. There were no significant differences in *Wolbachia* gene expression due to SINV infection. Both SINV+ and SINV- samples cluster closely together, and dimensions 1 and 2 are relatively short compared the *Drosophila* expression data.

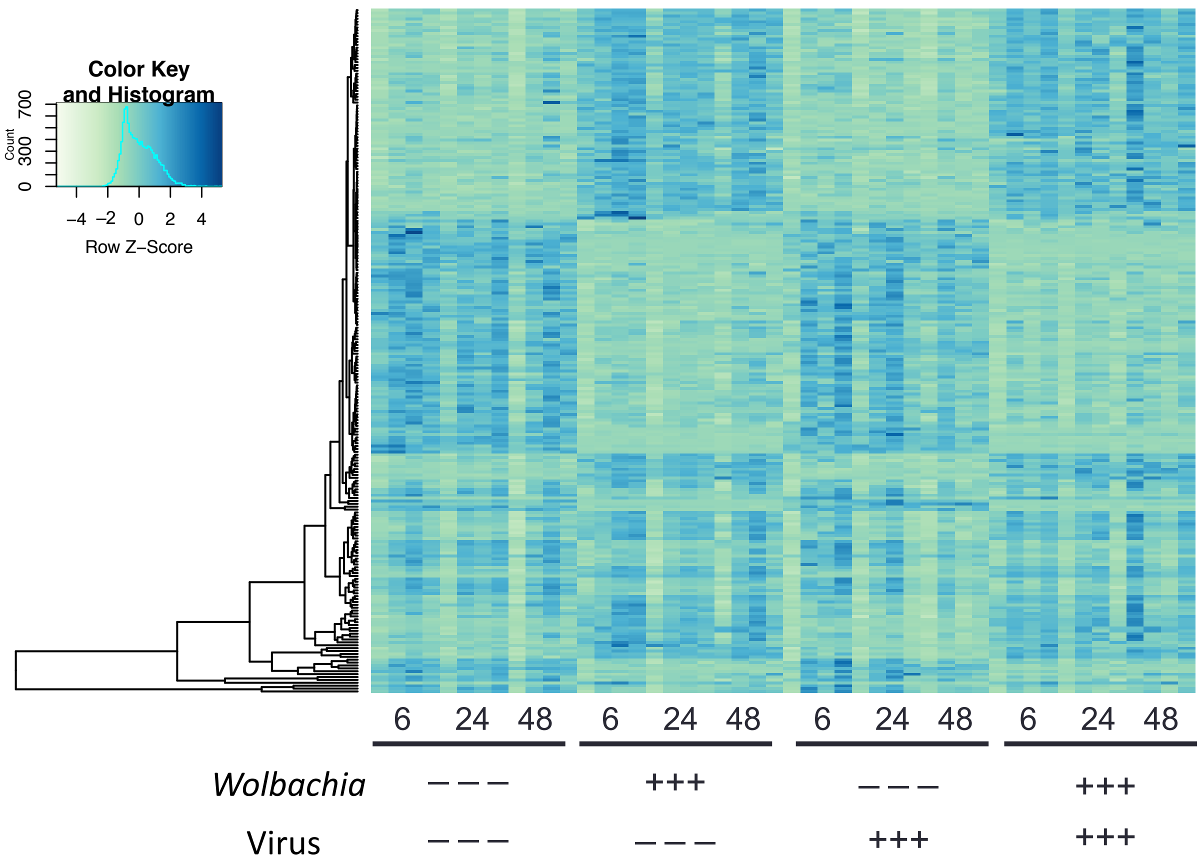


**Figure S2. Transcriptomic response to *Wolbachia* colonization**. Heatmap of the 237 genes significantly differentially expressed at the gene level, in response to *Wolbachia* colonization at an adjusted p-value of 0.05 and a fold change >2. *Wolbachia*-colonization, SINV-infection and timepoint are indicated under each set samples, with biological replicates adjacent to each other.

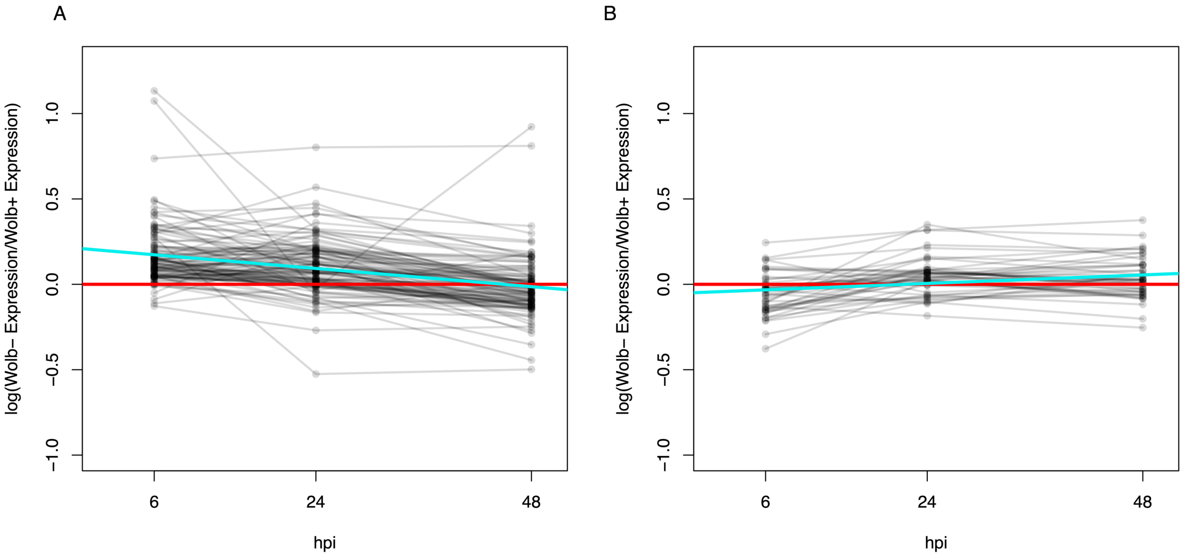


**Figure S3. *Wolbachia* colonization results in a muted response to virus infection.** At each time point and for each gene that was significantly differentially expressed due to SINV infection, the log of fold change in gene expression (based off TMM normalized values) was calculated for *Wolbachia*-uninfected samples injected with SINV (W-, SINV+), relative to *Wolbachia*-colonized samples injected with SINV (W+, SINV+). Values on the y-axis indicate how similar or dissimilar gene expression is, with a value of zero (red line) indicating the same level of expression in the W+ and W- samples for that gene. A larger value indicates the W- sample had higher expression for a given gene. Each grey line represents one differentially expressed "virus-responsive" gene. The blue line is a linear regression indicating whether or not the set of genes become more similar or more dissimilar in their expression between the W+ and W- samples over time. (A) Genes that are significantly upregulated in response to SINV. Positive values indicate a more exaggerated response in the W- sample. (B) Genes that are significantly downregulated in response to SINV. Negative values indicate a more exaggerated response in the W- sample.
